# Complete pangenome graph annotation with GrAnnoT and GrAPEx

**DOI:** 10.64898/2026.09.28.754982

**Authors:** Nina Marthe, Matthias Zytnicki, Tiziri Tamani, Francois Sabot

## Abstract

**Background:** Pangenome variations graphs (PVG) are promising structures to study diversity and reduce reference bias in genomics. However, it is not currently possible include complete annotation of the PVG. In a previous work, we introduced GrAnnoT, a tool for transferring the annotation from a reference genome to a PVG, and from a PVG to its embedded genomes. However, GrAnnoT was limited to transferring one annotation to the graph, and thus could not include all genes.

**Results:** We implemented new options to GrAnnoT, that can now handle annotations of multiple genomes, and integrate them in the graph. We also developed GrAPEx, a pipeline that identifies, extracts and annotates non-annotated regions in the graph. We tested those two tools to attest their performances. Our benchmarks revealed that both GrAnnoT and GrAPEx are reliable. GrAnnoT is now able to integrate correctly several annotations, increasing the amount of biological information available in the graph. Additionally, it offers useful options to study gene distribution in the PVG. GrAPEx allows rapid annotation of the whole PVG, with no reduction in the annotation quality.

**Conclusion:** Overall, the two tools presented here allow to annotate thoroughly the whole PVG, and integrate biological annotation to the pangenome. It also reduces the previous reference bias, where only one annotation (often the reference one) could be included in the graph. This constitutes a precious addition to the pangenomic toolkit.

## 1 Background

Advances in sequencing technologies and assembly methods have given access to a large number of new genomes in the past few years. In particular, multiple genomes for the same species can be produced at affordable cost [1–3]. These datasets enable intra-specific studies that compare whole genomes, and show that the use of a unique reference genome has limitations [4–6]. In particular, genomic sequences that are not on the reference genome are more difficult to study [5, 7]. However, these sequences harbour interesting information, and include functional elements linked to adaptation and evolution [8, 9]. Overall, using a single genome to represent a species biases many analysis, limits the study of the dispensable genome, and obstructs intra-specific diversity [10].

Pangenomes, encompassing information from several genomes from the same species or group [11], have been proposed to study such diversity and take into account regions absent from a single reference genome. They have been developed for many species, with applications in agronomy, medicine, evolution, etc [12–15]. These pangenomes can have different structures, from genes collection to whole genomes. In particular, the pangenome variation graph (PVG) [16] aims to represent a multiple genome alignment from various individuals. It includes the complete sequence from the genomes it represents, and allows to easily know what sequences are shared, and where are the differences. As it is often built from the alignment of complete genomes, it is well suited to detect structural variants, that are difficult to study in reference genomes [12]. In these graphs, nodes contain sequences and links represent adjacencies between the sequences. They are built by aligning whole genomes [17–19], and therefore include intergenic regions, in addition to gene-only sequences. This structure also informs on shared and different sequences, and is well suited to study eukaryotic pangenomes [12, 20].

As with the sequences present in genomes, the sequences included in this structure benefit from annotation, in order to add biological information and help interpret the variations found within the pangenome. In a previous work, we presented a first version of GrAnnoT [21], a tool developed to efficiently transfer an annotation from an embedded linear genome to a pangenome variation graph, and from the annotated PVG to its other embedded linear genomes. It projects annotations’ coordinates through the graph, and its informative outputs allow to review the transfers performed. We demonstrated that its performances are on par with Liftoff [22], the state-of-the-art annotation transfer tool for linear genomes, while being faster.

However, an important former limitation of GrAnnoT was that it could only integrate one annotation in the PVG. Therefore, any sequence not on the annotated genome would have no annotation. This restriction is impactful and introduces a strong reference bias. Indeed, one of the goals of using a pangenome is to include more sequences - and thus more genes - that are absent from the reference (and generally annotated) genome. To solve this limitation, we developed two extensions to GrAnnoT that allow to annotate the PVG more completely, or even in its entirety.

The first one is a modification of GrAnnoT itself that can now integrate several annotations in the PVG. That way, a larger part of the dispensable genome is annotated in the graph. To handle features shared between assemblies, GrAnnoT groups them into SynteGroups (groups of syntenic features), and can provide a presence-absence matrix based on these SynteGroups (SG). Since the PVG is built by aligning complete genomes, each gene’s sequence alignment is heavily dependant on the genomic context around it. Therefore, only genes with similar genomic context are aligned in the graph, and syntenic genes are grouped. At the opposite, approaches based on gene or protein sequence alignment do not take this context into account, and align regions that have similar sequence but not necessarily similar backgrounds. In consequence, these approaches group gene families, that include several related genes. GrAnnoT’s grouping, by relying on synteny, separates orthologues and paralogues, and groups alleles of the same gene. These allelic series enable the comparison of alleles, and the study of gene evolution within a species.

If not all genomes are annotated, some regions of the graph can still lack annotation information. To address this issue, the second extension, GrAPEx, provides annotations through the targeted use of Helixer [23] for the regions of the PVG that are not covered by an annotated genome, without any data needed other than the graph. This allows to annotate the whole graph without re-annotating known regions, thus reducing computation and runtime. The annotations produced by GrAPEx are then transferred in the PVG using GrAnnoT.

Together, these two extensions allow GrAnnoT to add biological information to the whole PVG and provide its full potential for studying the diversity it contains.

## 2 Results

### 2.1 Implementation

#### 2.1.1 GrAnnoT’s extension

The new version of GrAnnoT can integrate several annotations in the graph, and group features that are on overlapping positions in the graph (see Figure 1). This process (see Figure 2) is iterative: each annotation is loaded into the graph, and compared to the previous annotations already integrated. When a feature from the new annotation overlaps with a feature from the already-loaded annotations, they are considered similar. They are then compared to see if they are identical, in which case only one copy is conserved. Otherwise, they are simply grouped in the same SynteGroup (SG), that includes features that share nodes in the graph and thus have sequence similarity.

**Fig. 1.**
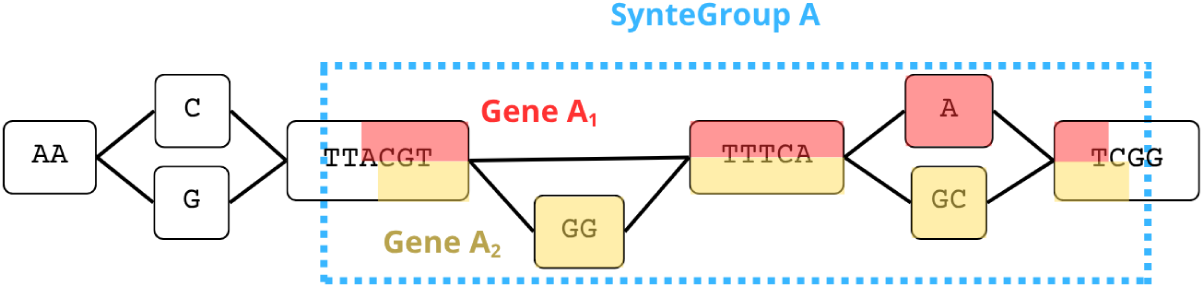
Similar annotations grouping in a SynteGroup. When two genes’ annotations overlap in the graph (*i.e.* when they share nodes), they are grouped in a SynteGroup.

**Fig. 2.**
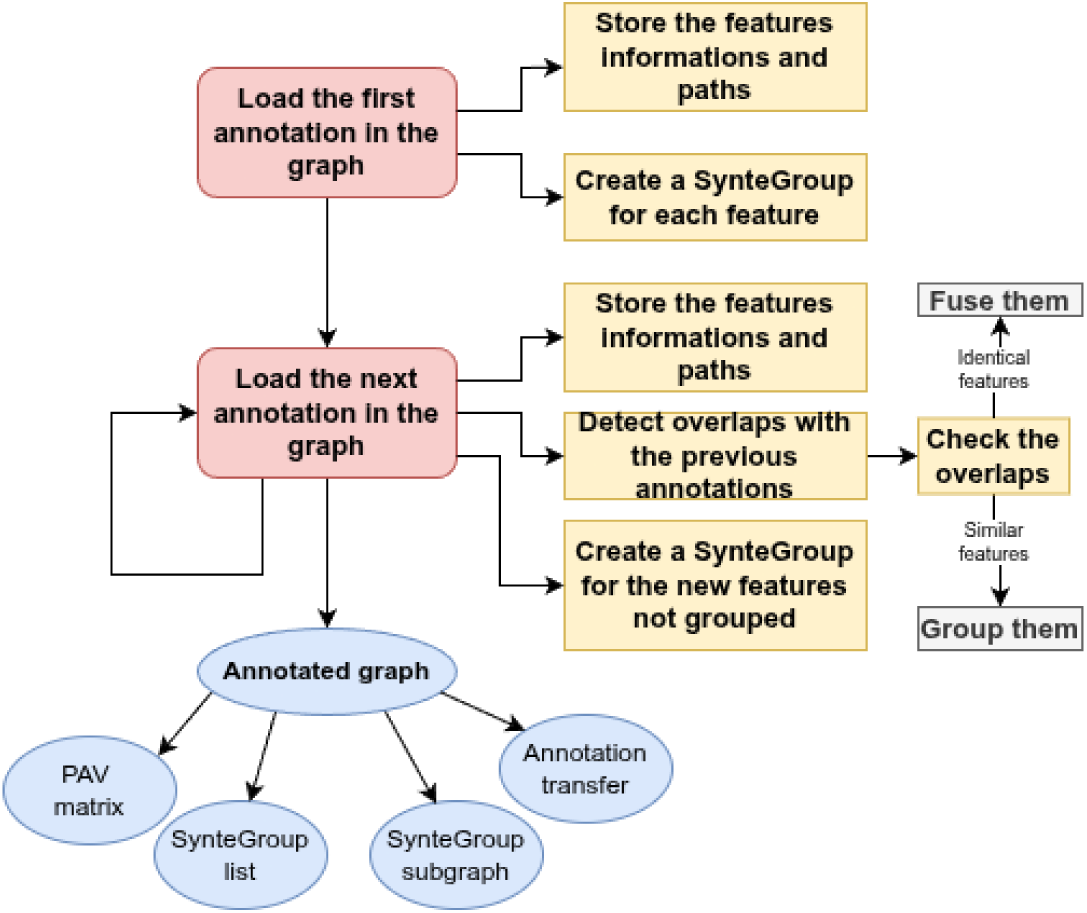
Multiple annotation transfer in the graph with GrAnnoT. To integrate multiple annotations in the graph, GrAnnoT loads them iteratively and detects the overlaps between genes of different annotations. Once the overlapping genes have been grouped, various outputs are available to describe the graph annotation.

Optionally, the parameter *overlap rate* can be used to filter the grouping of features. When it is used, when two features overlap, a similarity score is computed between them. This similarity score must reach the *overlap rate* value for the features to be grouped. The similarity is first evaluated between whole genes, then if the threshold is not reached, only the exonic nodes are considered (to negate the effects of intronic variations). The similarity between genes or exons is evaluated by computing a Jaccard index of the sequences (shared sequences length / total sequences length for the two genes or exon sets). If the *overlap rate* threshold is reached, the two genes are grouped. Since its default value is 0, by default all overlapping features are grouped (see section 2.2.1).

Once the overlaps are processed, features that do not overlap with any other one are put in their own SynteGroup.

After all the annotations are loaded in the graph and all the SynteGroups (or SG) are built, several outputs are available:

- Graph annotation with all features in GAF format (see [21] for details).
- Presence-absence matrix of the graph annotation, summarising the SG distribution of the features.
- Sub-graph extraction of a SG in GFA format, to visualise and compare the different versions of a feature.
- SG list, listing the features that form each SG.

The user can still transfer the graph annotation to target genomes. To avoid redundancy, GrAnnoT transfers only one feature per SG and selects the best copy. This selection is operated by finding the feature closest to the target genome (the one with the best coverage and sequence identity scores after transfer). If the graph annotation is transferred on target genomes, the presence-absence matrix includes the transfers performed.

#### 2.1.2 GrAPEx

When only a part of the genomes are annotated, regions outside of these genomes in the graph lack information. These regions can contain important genes, that risk being overlooked in favour of the annotated regions. To add annotation to these dispensable regions, it is possible to annotate the linear genomes that cover them. However, doing so would result in the re-annotation of known regions, producing redundant annotation.

In this context, GrAPEx was developed to extract and annotate only these non-covered regions. It uses Helixer [23] for *ab initio* annotation, and its process is described in Figure 3.

**Fig. 3.**
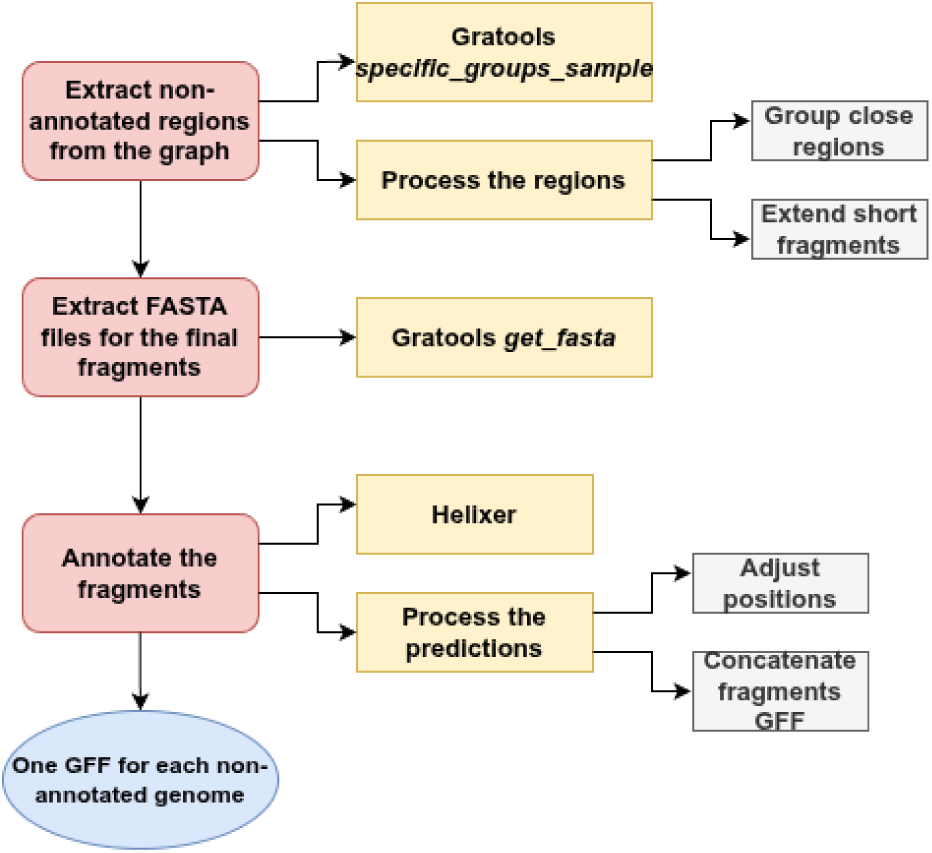
Annotation of non-covered regions with GrAPEx. GraTools is used to extract non-covered nodes from the graph. GrAPEx then processes these nodes to build fragments long enough. Their sequence are extracted from the graph using GraTools again, and the resulting fasta are given to Helixer. Finally, Helixer fragment annotations are processed to output genome annotations.

Firstly, GrAPEx uses GraTools [24], a PVG manipulation tool, to extract regions that are not covered by the annotated genomes (see section 5). Once fragments larger that 1kb are extracted for each non-annotated genome, they are processed to reach a minimum fragment length threshold (100kb by default), to guarantee a better annotation quality with Helixer. Indeed, from our tests, Helixer performs worse on small sequences. To increase the length of the fragments, they are first aggregated with nearby fragments. Then, the remaining fragments that still don’t reach the minimum length threshold are extended (see Figure 4). After the fragments have been lengthened, GrAPEx calls Helixer to produce *ab initio* annotations on their sequences. The resulting GFFs are then parsed to keep only the annotations corresponding to the targeted regions. Finally, they are merged and the annotations’ positions are adjusted to produce a single GFF for each of the non-annotated samples. These final annotations can be given as input to GrAnnoT, to be transferred into the graph.

**Fig. 4.**
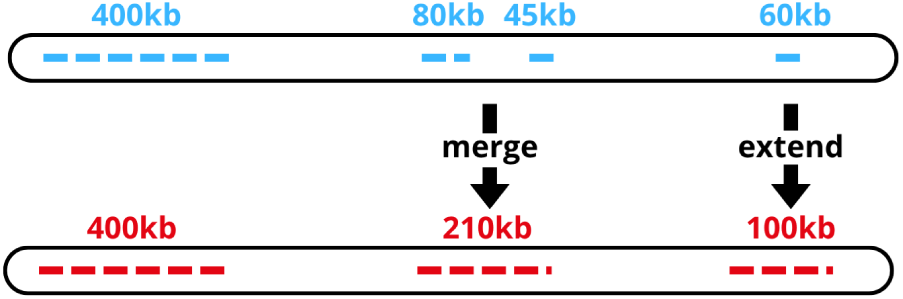
Fragment size adjustment prior to Helixer annotation. To reach a minimum fragment length of 100 kb, GrAPEx first merges nearby fragments, and then extends those that still do not reach the threshold.

### 2.2 GrAnnoT benchmarks

#### 2.2.1 BUSCO genes grouping

To ensure that we grouped correctly the genes shared between the annotations, we built a dataset based on single-copy BUSCO genes [25] in rice genomes (see section 5). In this dataset, each genome has an annotation comprising its BUSCO genes, including only shared, conserved, and single-copy genes [25] and removing duplicated ones (in individual genomes). That way, we know how the genes are supposed to be grouped: each SynteGroup is expected to include the different copies of one unique BUSCO gene, with maximum one copy per embedded genome.

The annotations of this dataset were transferred on the graph and grouped in SynteGroups using GrAnnoT. We then verified that all the copies of each BUSCO gene were grouped in the same SynteGroup, and not split in the graph. We repeated this analysis using different values for the parameter *overlap rate* (described in subsection 2.1.1), to evaluate its impact in this controlled setting.

As we can see in Figure 5, with an *overlap rate* of 0, 98.4% of BUSCO genes are grouped in only one SynteGroup. This confirms that the graph aligned correctly these annotated regions, and that GrAnnoT managed to detect the overlap.

**Fig. 5.**
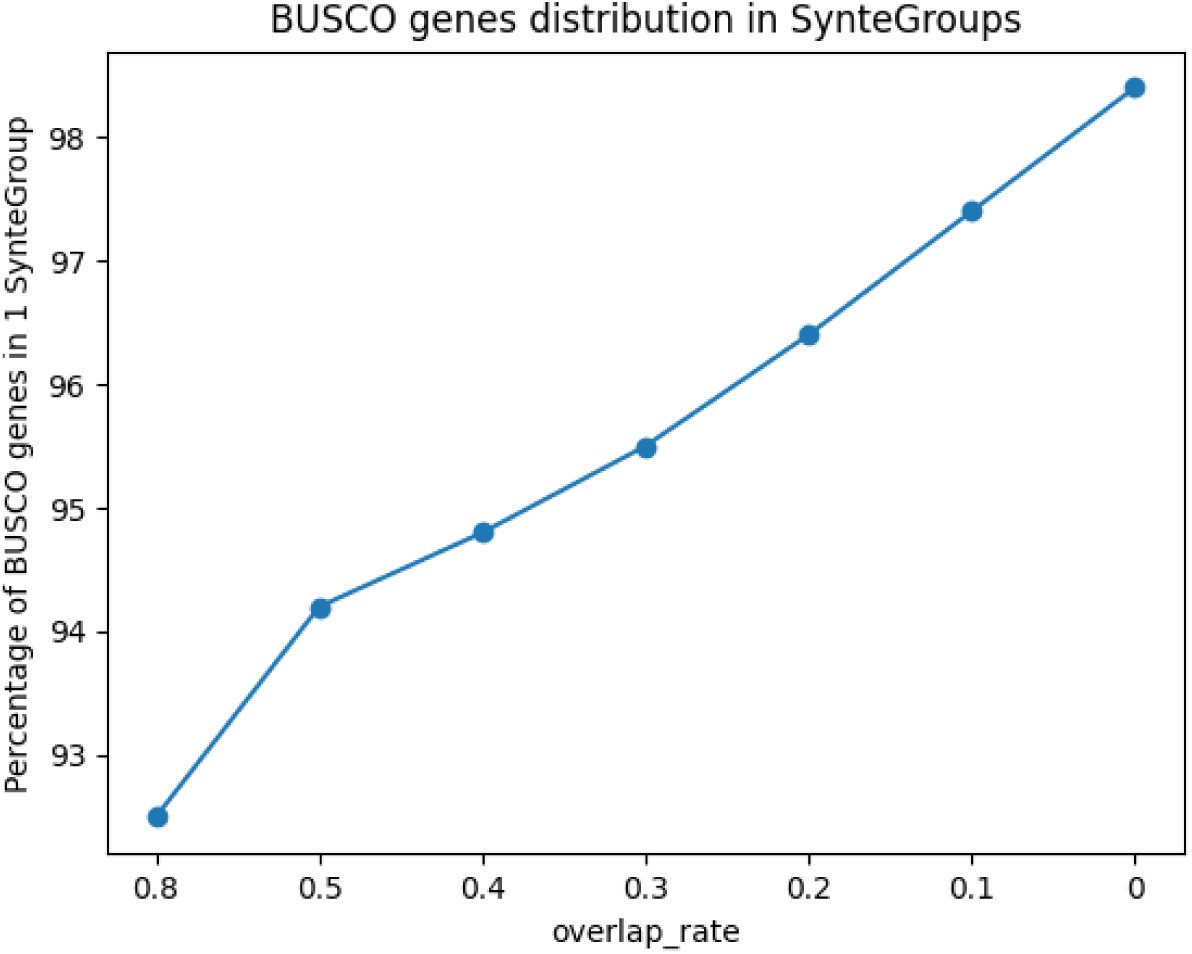
BUSCO genes distribution in SynteGroups. Percentage of BUSCO genes in exactly one SynteGroup, with varying values for the parameter *overlap rate*. The higher percentage is reached for *overlap rate* at 0.

We can also see the importance of the parameter *overlap rate*. This parameter corresponds to the minimum overlap score between two features to group them (see part 2.1.1 for details). As showed in Figure 5, an *overlap rate* too high looses relevant grouping. For our dataset, the best grouping occurs when any overlap is accepted to group two features, with no minimum score to reach.

Overall, depending on the data, the species, and the scientific question, this *overlap rate* parameter could be adjusted by the user.

#### 2.2.2 Comparison to an existing method : *get pangenes* on the Panoryza dataset

In a paper from 2026, B. Contreras-Moreira, E. Sharma, S. Saraf *et al.* applied the method *get pangenes* [26] to a Asian rice pangenome dataset embedding 16 platinum individual rice genomes, to combine individual annotations from these genomes based on synteny and whole genome alignments [27].

By transferring several annotations in a pangenome graph, GrAnnoT can group them in a similar manner. Compared to *get pangenes*, the approaches have differences, but both rely on synteny and whole-genome alignment. This common characteristic allows to compare them more easily than other methods that rely on gene or protein alignment against each other or against a genome [28–30], and therefore group gene families instead of allelic series. Therefore, we chose to reproduce some analysis from the *get pangenes* method with GrAnnoT SynteGroups.

The Figure 6 compares the data from Panoryza [27], using the method *get pangenes* [26], and the current version of GrAnnoT, using the same figure structure as the Figure 1.A from Panoryza paper. We can see that the two approaches are similar: in both cases, the most prevalent occupancies are 1 (singletons) and 16 (core pangenes/SG). For the core pangenes/SG, the two approaches gave close results, with around 25k core pangenes and SG. For the singletons, GrAnnoT displays less SynteGroups, and has more elements shared by 2-15 genomes compared to the *get pangenes* method (∽18k *vs* ∽9k).

**Fig. 6.**
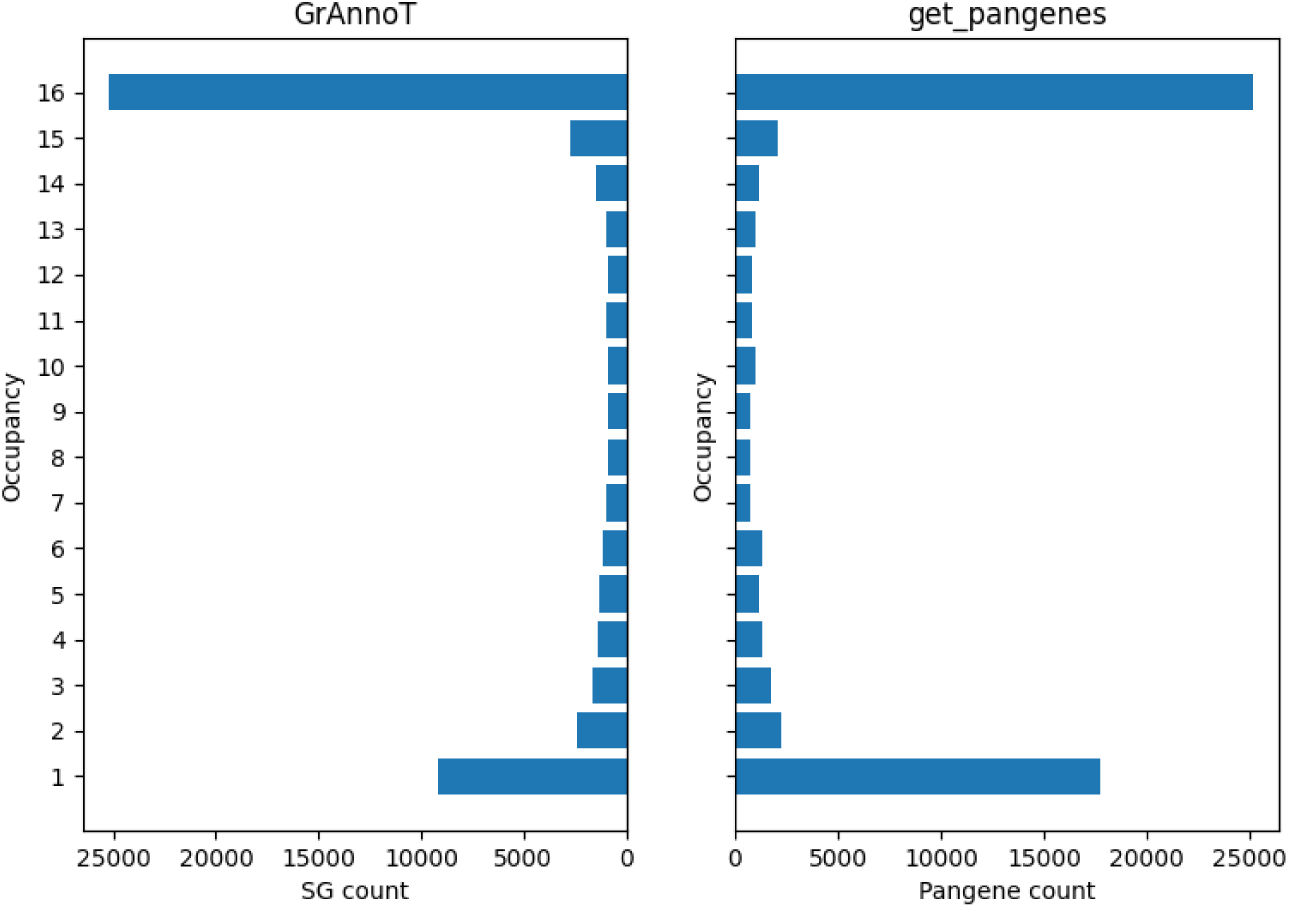
Pangene/SynteGroup occupancy. The number of elements (SG or pangene) per occupancy (number of genomes involved) is represented. The right panel represents the pangenes, and the left one the SynteGroups.

On Figure 7 is represented the core gene growth curve for the two methods, similar to the Figure 1.B from the paper Panoryza. There, we see that the shapes of the curves are similar, with the number of core pangenes/SG declining with more genomes added. As seen in Figure 6, both methods give ∽25k core elements (pangenes or SG).

**Fig. 7.**
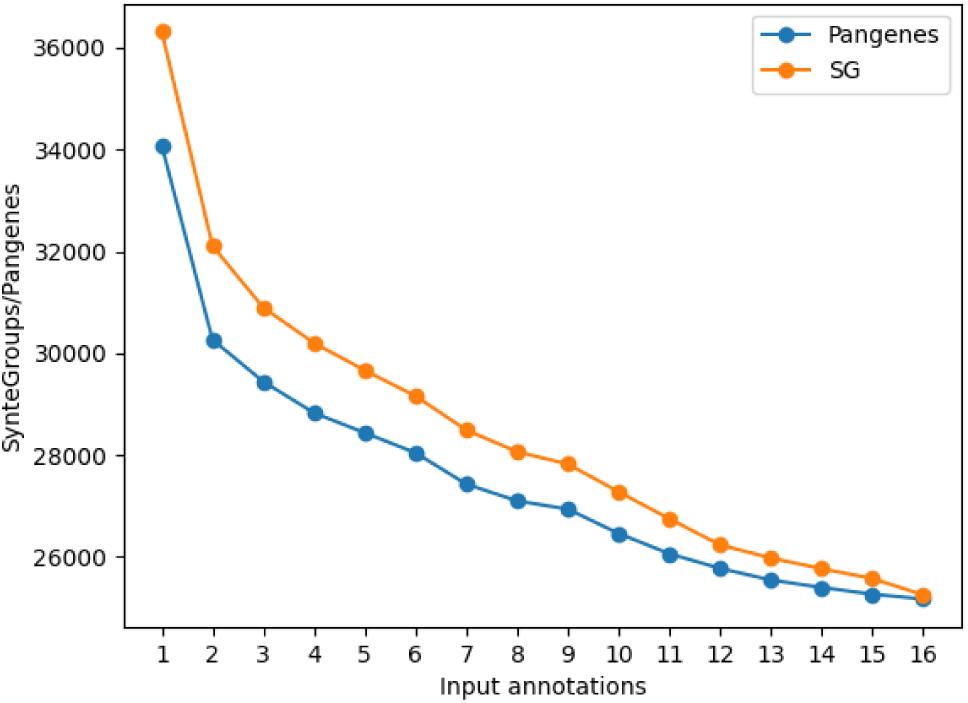
Core genes growth. Number of pangenes (in blue) and SG (in orange) shared by all genomes for 1-16 genomes.

The Figure 8 illustrates the singleton numbers, similarly to the Figure 1.C from Panoryza. As with Figure 6, GrAnnoT produces less singletons than *get pangenes*, although with the same order of magnitude for the genomes (between ∽400 and ∽1000 for most genomes in both cases). Their distribution is also different between the two methods, as the genomes with the more or the less singletons are not shared.

**Fig. 8.**
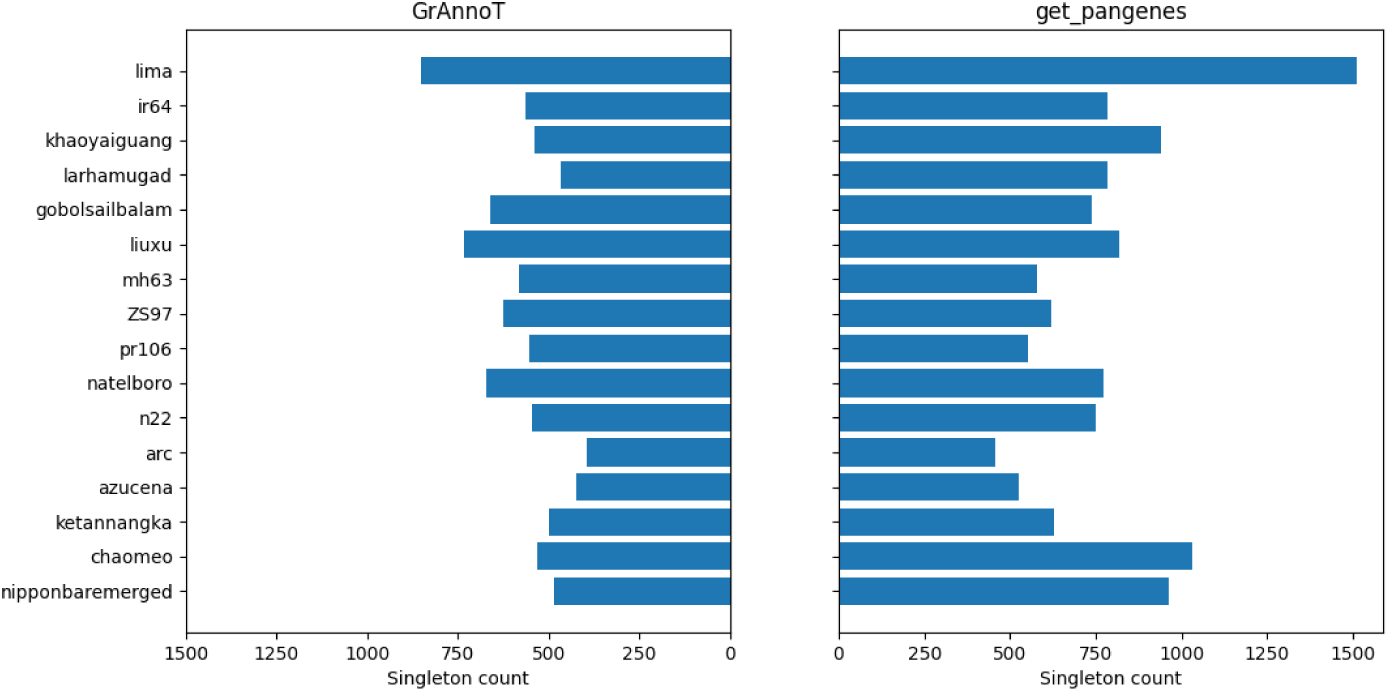
Singleton distribution. Number of specific pangenes (right panel) and SG (left panel) for each genome.

Finally, Figure 9 highlights the population structure based on pangene/SG sharing, similarly to the Figure 1.D from the paper Panoryza. Both approaches find that the Nipponbare annotation is very different from the rest. This is expected, since the Nipponbare annotation used was built by merging 3 different annotations. It has a higher gene count than the other annotations, and thus shares a lower portion with the other genomes (see the original Panoryza paper [27]). The genomes clustering based on SG sharing appears to correspond better to the population structure than the clustering based on pangenes, with genomes from the same rice sub-families closer in the phylogenic tree. Apart from Nipponbare, the clustering matches exactly the population structure. When Nipponbare annotation is filtered to keep only the genes from the same source as the other genomes (and from only one annotation), Nipponbare is grouped correctly with the other japonica genomes (see Supplementary Figure A1).

**Fig. 9.**
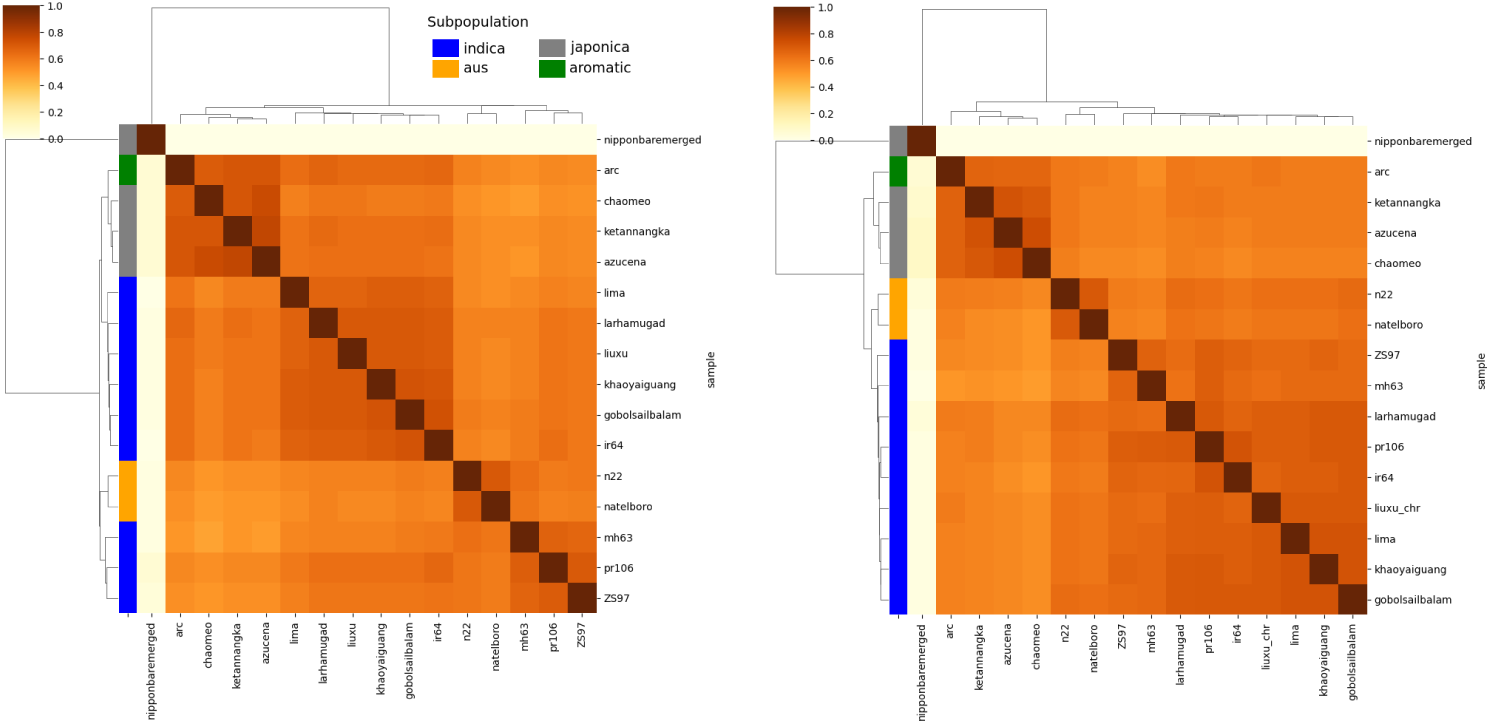
Pangene/SG cluster overlap. Matrix of shared pangenes/SG between genomes. The figure on the left represents the pangenes, and the figure on the right represents the SynteGroups.

Overall, this comparison shows that the pangene clustering from *get pangenes* has similar tendencies compared to GrAnnoT. Both approaches find the same number of core elements (∽25k, Figures 6 and 7) but GrAnnoT finds less singletons, and the population structure is better represented with the SG compared to the pangenes (Figure 9).

#### 2.2.3 Graph annotation growth

Using GrAnnoT, when a supplementary annotation is added in the graph, any gene shared with the previous annotations is grouped in an existing SynteGroup (or SG). Therefore, as new annotations are added in the graph, the SynteGroups should include more genes, and the total number of SynteGroups should not increase as fast as the total number of genes. Additionally, when two genes are identical, only one copy is kept in the graph. Thus, some genes will be deleted and the number of genes in the graph should be lower that the total number of input genes.

To quantify this, we counted the number of SG, the number of genes in the graph, and the total number of genes in the input annotations, with 1 to 16 annotations transferred in the Asian rice graph (see section 5 for details on the data used).

We can see on Figure 10 that the total gene count is slightly reduced in the graph due to identical annotations. This behaviour allows to save space and reduce redundancy in the graph annotation. In this example, it reduced the number of individual genes by 16%. Additionally, as expected, the gene count in the graph increases much faster than the SG count. This reduction is much more important than the previous one (86.4%). This demonstrates that while genomes share genes, they usually have different versions or alleles of these genes, and intra-genic variations are frequent. However, the different gene versions in the SG can partly be explained by discrepancies in the genomes’ annotations.

**Fig. 10.**
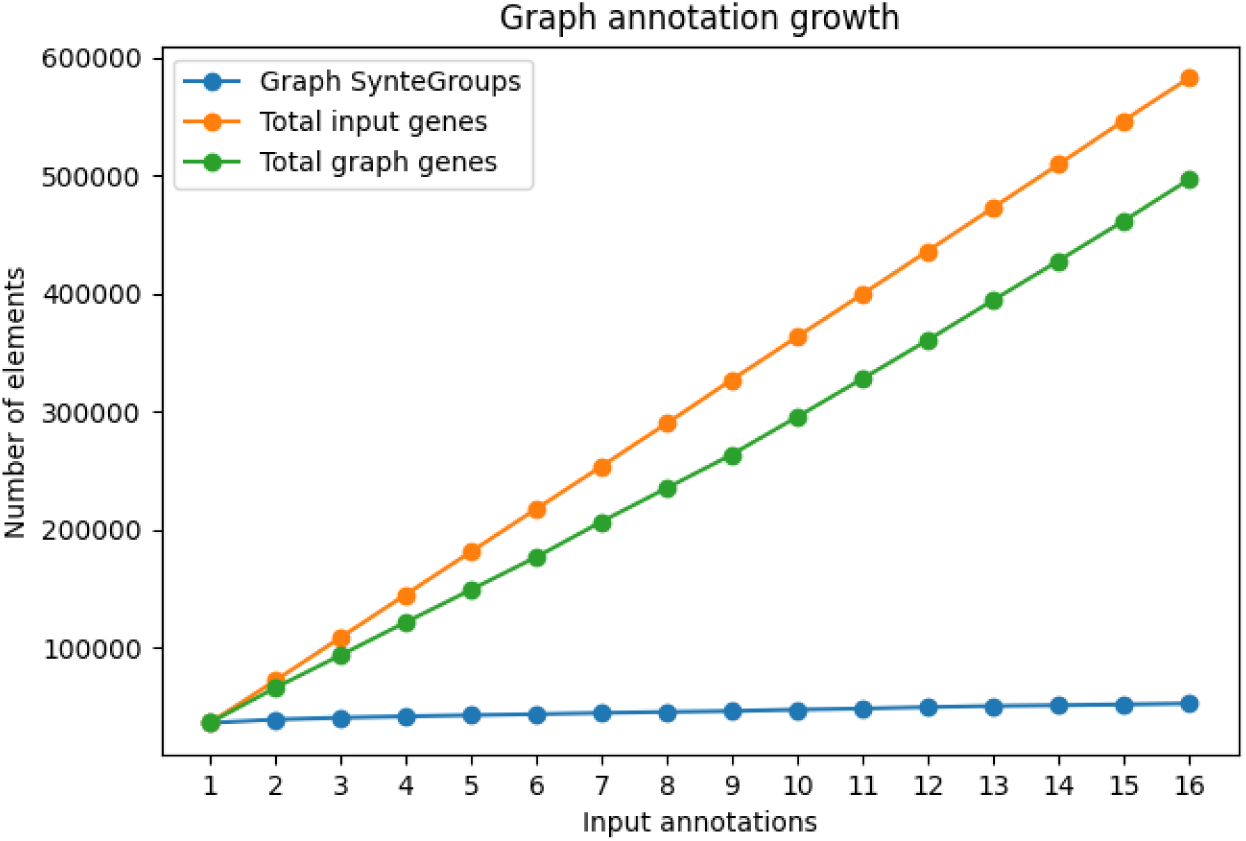
Graph annotation growth. Comparison of the number of input genes, graph genes, and graph SynteGroups with 1-16 input annotations.

Overall, this shows that GrAnnoT is indeed able to detect identical or similar features, and group them to better represent the gene distribution within the pangenome.

#### 2.2.4 Target genome annotation transfer growth

By transferring several annotations in the graph, we increase the quantity of dispensable regions annotated. Thus, the annotation transferred on a target genome is more complete when several annotations are integrated in the graph.

We demonstrated this gain in target genome annotation by transferring the graph annotation based on a growing number of genomes (see Figure 11). The graph annotation and the SynteGroups were computed by transferring in the graph 1, 2, 3, …, up to 16 annotations. The graph annotation was then transferred on the last genome, Nipponbare. Finally, we counted the number of genes transferred on Nipponbare.

**Fig. 11.**
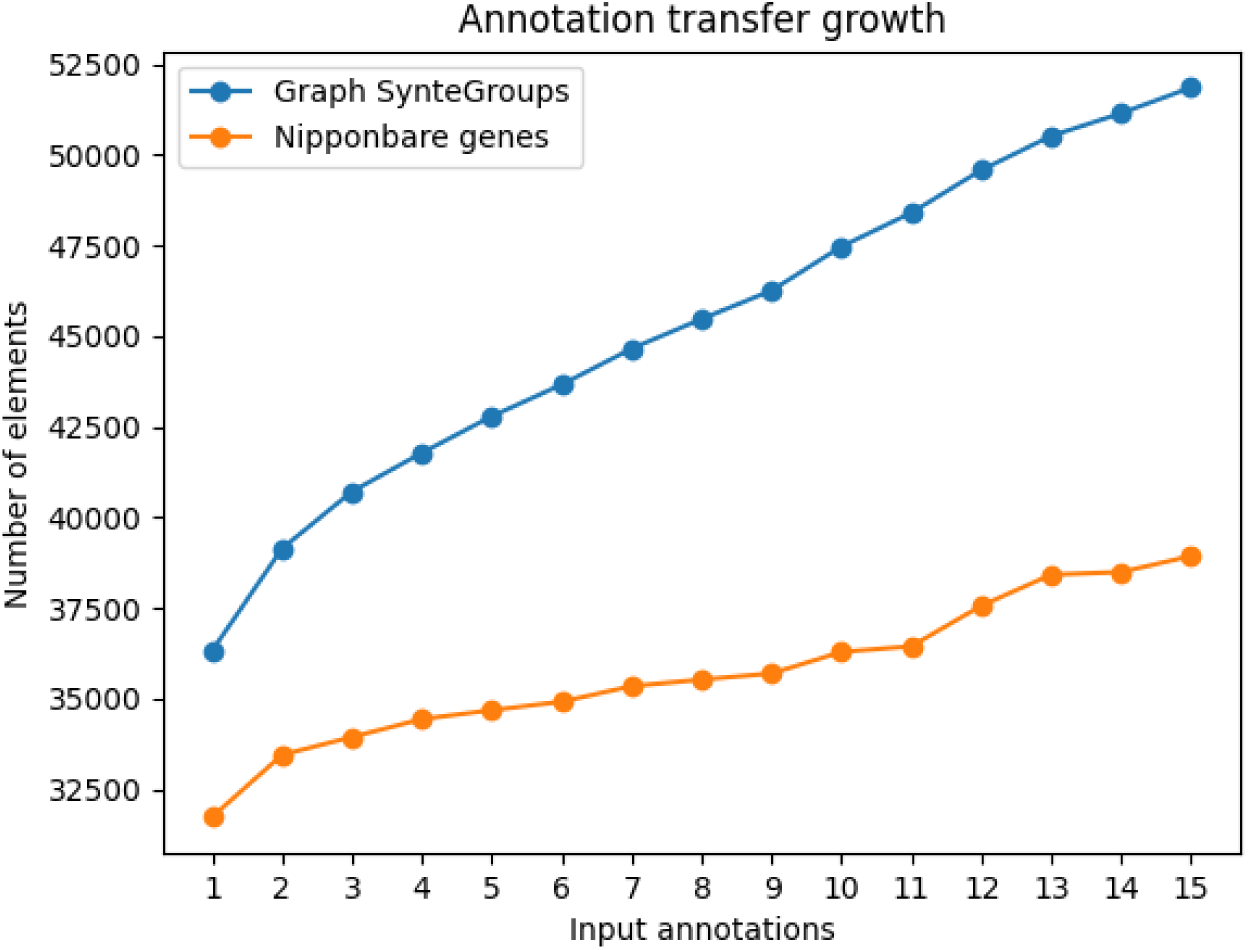
Annotation transfer growth. Comparison of the number of SynteGroups in the graph and genes transferred from the graph to genome Nipponbare, with 1-16 input annotations.

As expected, the more we add annotations in the graph, the more SynteGroups there are, and the more annotations are transferred on the last genome.

After the first annotation, the curves on the figure are overall linear, for the SynteGroup especially. The number of new SynteGroup per genome annotation added does not seem to decrease : after the first genome’s annotation, each new genome annotation adds a similar amount of new genes in the graph. This means that the pangenome annotation is not saturated, suggesting an open pangenome [31], or that the gene diversity of Asian rice is not represented in its entirety in this pangenome.

### 2.3 GrAPEx benchmarks

GrAPEx was developed to annotate graph regions that are not covered by annotated genomes, without other data than the graph. That way, the graph annotation is less redundant and the total quantity of sequence to annotate is reduced for faster runtime.

#### 2.3.1 Reduction of the amount of sequence to annotate

To observe more precisely how much GrAPEx reduces the amount of sequence to annotate, we compared the cumulated length of sequence annotated using GrAPEx versus whole genome annotation. We also measured the time needed for these two approaches.

In table 1, we tested GrAPEx on a pangenome graph with 16 genomes in total, with only one annotated. We can see that using GrAPEx allows to reduce the total sequence to annotate by 73.8% (from 5.92 Gbp to 1.55 Gbp), and the runtime by 45.7% (from ∽38h11 to ∽20h45).

**Table 1.** Pangenome annotation with Helixer.

| Annotated sequence | Annotated sequence length | Annotation and transfer runtime | Max memory |
| --- | --- | --- | --- |
| GrAPEx fragments | 1.55 Gbp | 20h45 | 36.24 Mb |
| Full genomes | 5.92 Gbp | 38h11 | 31 Mb |

#### 2.3.2 Increase in the annotated portion of the graph

By augmenting the graph annotation, GrAPEx allows to transfer more dispensable annotations on target genomes. Similarly to Figure 10 in subsection 2.2.4, we verified this by comparing the number of SG in the graph and the number of features transferred on a target genome, with one whole genome annotation (here from genome Larhamugad) and including or not GrAPEx annotation of the excluded regions.

Table 2 shows that as expected, GrAPEx completes the graph annotation by adding new genes and SG. Integrating GrAPEx annotations adds a similar amount of SG compared to integrating whole genome original annotations (53548 *vs* 52708).

**Table 2.** Pangenome annotation with Helixer.

| Condition | Larhamugad | Larhamugad+GrAPEx | Larhamugad+15 genomes |
| --- | --- | --- | --- |
| SynteGroups | 36316 | 53548 | 52708 |

For the graph annotation transferred on genome Nipponbare, table 3 shows that adding GrAPEx predictions to the graph enriches the Nipponbare annotation transfer as expected. The total number of genes in Nipponbare using GrAPEx annotations is closer to the expected amount than when only one annotation is used.

**Table 3.** Pangenome annotation with Helixer.

| Condition | Transferred from Larhamugad | Transferred from Larhamugad+GrAPEx | Nipponbare annotation |
| --- | --- | --- | --- |
| Nipponbare genes | 31736 | 35913 | 36212 |

#### 2.3.3 Impact of fragmentation on annotation quality

The runtime advantage of GrAPEx demonstrated in subsection 2.3.1 stems from the fact that it only annotates fragments of the non-annotated genomes. However, the *ab initio* annotation tool that it uses, Helixer [23], has lower performance on small genomes or sequences. When developing GrAPEx, we thoroughly tested the length of the fragments, to guarantee a sufficient annotation quality. This lead us to add a length threshold parameter (100kb by default), and the fragments not reaching it are extended (see part 2.1.2).

Here, we verify that the fragmentation of the sequences extracted by GrAPEx does not impact their annotation by Helixer, using the first extracted genome, ZS97.

We first tested that the fragments extracted by GrAPEx are indeed 100kb long (see Supplementary Figure C2). Then, we compared the annotation done on the fragments to the annotation done on the complete genome. Firstly, we checked that both protocols generated a similar amount of predictions. On whole genomes, Helixer predicted 3588 genes, and on GrAPEx fragments, 3149 genes. Then, the predictions themselves were compared on Figure 12.

**Fig. 12.**
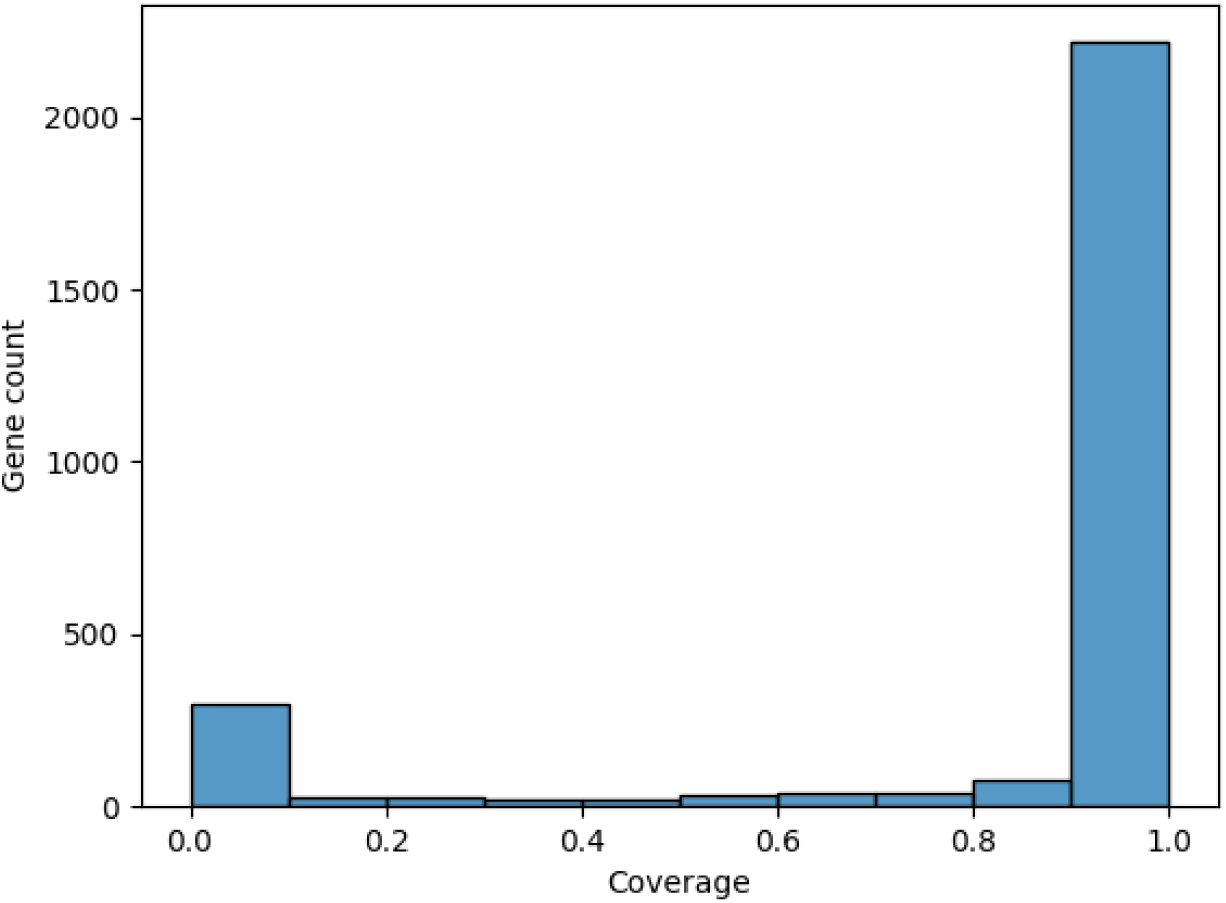
Helixer prediction comparison. Comparison between Helixer predictions on GrAPEx fragments *vs* on whole genome. The coverage corresponds to the portion of each gene from the fragments that overlaps with a gene from the whole genome.

We can see that annotating fragments or whole genome with Helixer changes moderately the positions of the annotated features. Around 80% of the features annotated on GrAPEx’s fragments overlap completely with features annotated on the whole genome. This demonstrates that while GrAPEx fragmentation of the genomic sequence to annotate reduces the amount of prediction, it has little impact on the positions of the predicted genes, thus on the quality of the annotation.

## 3 Discussion

Together, these two methods constitute precious additions to GrAnnoT to generate and handle complete pangenome graph annotation. That way, the pangenome variation graph structure can include biological information, that brings meaning to the variations it represents. It allows to have in the same structure whole genome sequences, small and large variations between the genomes, gene distribution in the pangenome, alleles collections, and intra-genic variations, without having to return to linear genomes. Additionally, it reduces GrAnnoT’s previous bias linked to the choice of the only annotation to be transferred. The annotated genome is often the reference one, and the two methods presented in this paper contribute to reduce reference bias, at the annotation level.

### 3.1 GrAnnoT-SynteGroups

GrAnnoT’s ability to aggregate several annotations leads to the concept of SynteGroups. They include syntenic features, with similar sequences. When the annotated elements are genes, they constitute an allelic series, with the different copies of a gene grouped in the same structure. Therefore, SG offer a way to separate orthologs and paralogs, and study intra-genic diversity and allele distribution within a pangenome.

With the benchmarks presented in this paper, we showed that GrAnnoT produces relevant SG clustering.

We first verified that the gene grouping was valid using BUSCO genes. As showed in subsection 2.2.1, these genes were generally correctly grouped. With the different *overlap rate* values we tested, between 7.5 and 1.6% of BUSCO genes were split between two or more SG. The best grouping occurred with *overlap rate* at 0, so we kept this value for all analysis with the same dataset.

When building the BUSCO dataset, we only had access to the positions of the genes, without any information on mRNA, exons, UTR, or CDS. This gene structure can help group similar genes, and could have improved the grouping of BUSCO genes if available. Indeed, when the overlap between two genes is insufficient, their exons are compared (see part 2.1.1). If the overlap between the two exon sets is sufficient, the genes are grouped. That way, differences in gene sequence due to intronic insertions are ignored. Thus, annotations with complete gene structures are easier to appropriately group, and if we had access to the BUSCO genes’ structure, we may have grouped them better with an *overlap rate>*0.

Next, the comparison to Panoryza paper showed good consistency overall. The only differences are that GrAnnoT’s method tends to group more genes and produce less singletons (see Figures 6 and 7), and its grouping follows better the population structure (see Figure 9). These comparisons demonstrate that GrAnnoT’s SG grouping is relevant, and allows to study PVG annotation efficiently.

As showed in subsection 2.2.3, SG grouping drastically reduces the amount of elements considered. Despite this grouping, all individual genes are conserved (except identical copies), and the user can always get back the entirety of their input data. However, by grouping genes into SG, the gene distribution in the pangenome is easier to see, with the SG presence-absence matrix for instance. Also, when transferring the graph annotation to a target genome, only one copy of each SG is transferred (see subsection 2.1.1), and the resulting annotation is much lighter and easier to manipulate for downstream analysis. For users interested in the whole pangenome annotation, including the different versions of each SG, the GAF output shows each copy. A subgraph of a SG can also be outputted, and visualized with tools like Bandage [32] to see in details the variations in the SG.

Finally, with more annotations and SynteGroups in the graph, the transfer to a non-annotated target genome is more complete, as seen in subsection 2.2.4. After transferring 15 annotations on the graph, the number of genes transferred on target genome Nipponbare is close to the expected number (38940 transferred for 36212 expected, so 7.5% more).

Overall, GrAnnoT’s integration of several annotations appears efficient. It informs on the population gene distribution and structure, and facilitates the study of a redundant dataset. SynteGroups construction seems relevant, and effectively groups related features.

The parameter controlling the SG construction, *overlap rate*, was set to 0 for the PVG used in this paper. The BUSCO dataset allowed us to determine the best value. However, the filtering of the overlaps could still be improved. Other methods could be used to evaluate the similarity between two annotations, like comparing protein sequences between overlapping coding genes. This would be more focussed on gene functionality and less sensitive to mutations. The evaluation of overlaps could also be more flexible, by letting the user choose different thresholds for the different overlap evaluations (whole gene vs only exons), or include other methods to evaluate overlapping genes.

In the meantime, GrAnnoT’s different outputs allow to review the SG grouping (SynteGroup list, PAV matrix, SG subgraph, graph annotation in GAF format). It can help the user to evaluate the relevance of the results, understand the results given by GrAnnoT, and adjust the *overlap rate* parameter if needed.

As any method, GrAnnoT results depend heavily on the input data that is provided, here the pangenome graph and the annotations. The overlap between the annotations and the transfers rely on the alignment between genomes that the graph represents, as GrAnnoT does not perform any alignment itself. As seen in GrAnnoT’s first paper [21], non-syntenic features tend to be poorly transferred, because of the original alignment that constitutes the PVG. As alignment and graph construction methods will get better, GrAnnoT will be able to handle more cases.

GrAnnoT is also dependant on its input annotations. Genes can be split or fused during the annotation process, and predictions can introduce artifacts. These annotations can disrupt the grouping, and errors can be spread in datasets when transferred to other genomes. In such cases, GrAnnoT informative output can be explored to see the annotations in the pangenome graph. Since all the input data is conserved, it is always possible to understand why GrAnnoT grouped or not two genes. For instance, the SynteGroup list shows for each SG what gene from what genome it contains; the GAF annotation of the graph shows the path of each feature, and can be used to see the overlap between two features in the graph.

Given GrAnnoT’s reliance to its input data, we could implement options to make it more flexible, to allow the user to adapt its behaviour to the characteristics of its data.

### 3.2 GrAPEx

The benchmarks conducted on GrAPEx demonstrated that it effectively reduces the amount of sequence to annotate and the annotation time, while providing annotation for all the regions of the graph. It was able to recover expected dispensable genes (see Appendix B), and proved its value. Therefore, it is a valid tool to annotate an entire pangenome with limited resources, when not all genomes are annotated.

We chose Helixer to perform the *ab initio* annotations. This tool offers acceptable predictions, and does not need any input data other than the sequence to annotate. No training or fine-tuning is required, and it is relatively fast [23]. The context in which Helixer is used for PVG annotation (*i.e.* on genome fragments) does not appear to impair significantly its performance. However, as any *ab initio* annotation tool, its predictions are fallible and should be carefully checked before subsequent analysis.

In the future, we would like to include other annotations tools to GrAPEx. It is already somewhat possible, since each GrAPEx step can be run separately. Thus, the user can retrieve the fragments extracted by GrAPEx, annotate them with the method of its choice, and then give them back to GrAPEx for the post processing steps. However GrAPEx was not conceived in that spirit, and changing the annotation method will be made easier in future developments.

## 4 Conclusion

In conclusion, GrAnnoT and GrAPEx offer to annotate completely a pangenome graph, thus integrating biological information in this structure. This allows to explore it more efficiently, and include more diversity in studies.

In this article, we presented these two methods, and tested their results. We demonstrated that they are valid and efficient, with few limitations.

We believe these tools can be help considerably people working with pangenome graphs.

## 5 Methods

All commands and scripts used to produce the data and the results are available in GrAnnoT repository on the IRD forge (https://forge.ird.fr/diade/dynadiv/grannot).

For all transfers with GrAnnoT, the parameter *overlap rate* was set at 0, except when stated otherwise (for the BUSCO benchmark). All other parameters were left at their default value.

### Data

For all benchmarks, the Panoryza pangenome graph was used. This graph contains the 16 Asian rice genomes described and used in the paper Panoryza [27]. It was built with minigraph-cactus [18], using 12 cores, the option *–gfa full*, and genome Larhamugad as a reference. The reference genome was chosen to minimize its average mash distance [33] to the other genomes, therefore choosing the barycentre of the pangenome diversity.

The annotations used also come from the paper Panoryza [27]. They have been adapted to GrAnnoT, since the GFF did not meet all requirement for our approach.

For the comparison with the results from the paper Panoryza [27], we used their Supplemental Table S1 (see details in 5).

### GrAPEx’s implementation

To extract non-covered regions from the graph, GrAPEx uses GraTool’s option *specific groups sample*. This option is run iteratively on all the non-annotated genomes. For each genome, all of its graph nodes that are not shared with either an annotated genome or an already extracted genome are retrieved.

Since the sequences are extracted genome by genome, they correspond to real genomic sequences and not chimeras built from the graph. Additionally, this approach does not require graph traversal, which is a difficult and costly operation.

After having retrieved all the non covered nodes per genome, adjacent node positions are joined using *bedtools merge* with the option *-d 0*. Then, the joined fragments are filtered to keep only those larger than 1 kb, to exclude the smaller insertions. Next, the fragments are joined again to raise the size of the sequences to annotate. This time, if two fragments are closer than 150 kb, their positions are joined, using again *bedtools intersect* with the option *-d 150000*. After joining the fragments for each genome, joined fragments still smaller than 100 kb are extended in 5’ and 3’ (if possible), to reach the minimum length of 100 kb. The fragments that were larger than 100 kb are extended of 10 kb on each extremity, to include enough context for the genes placed at the extremity of a fragment.

Once these positions are computed, the corresponding fasta files are extracted from the graph using GraTool’s *get fasta* option. The fasta files are then given to Helixer [23] for *ab initio* annotation. Then, the GFF are parsed to only keep genes that were in the original extracted positions (corresponding to non-annotated nodes in the graph). Finally, the positions of the genes are concatenated and adjusted to correspond to the whole genome, and not their fragment. In the end, one GFF is produced for each non-annotated genome.

### GrAnnoT-SG benchmarks

#### BUSCO genes grouping

BUSCO [25] was run on all 16 genomes of the Panoryza dataset, using the lineage *Poales*. Among BUSCO’s outputs, the table describing the genes found in the genome (*full table.tsv*) was parsed to get the genes’ positions. GFF files were built with these genes, and include their positions and their ID. Duplicated genes were excluded, to simplify the dataset.

The 16 BUSCO annotations were transferred on the graph using different *overlap rate* values, and the list of SynteGroups was outputted for each *overlap rate* value. These SG lists were then parsed to analyse how the BUSCO genes were grouped by GrAnnoT.

#### Comparison to *get pangenes*

GrAnnoT was run with the Panoryza data to reproduce the Figure 1 from the paper Panoryza [27]. For Figures 6 8 9, all 16 annotations were transferred in the graph, and the SG list was outputted and parsed to extract the informations needed for the figures. For Figure 7, 1 to 16 annotations were transferred in the graph with a fixed order. The first annotation transferred is Larhamugad’s one, and the other are integrated in growing mash distance [33] order. This order is the same as the one used for the graph construction with minigraph-cactus. For this figure, the PAV matrix was outputted and parsed.

To reproduce Panoryza figures, the Supplemental Table S1 from the paper Panoryza [27] was parsed. For Figure 7, we analysed the pangenes to simulate the integration of 1 to 16 annotations, keeping the same order as with GrAnnoT (see above). When appropriated, pangenes were filtered to exclude those with only a gene from Nipponbare’s MSU annotation, as was done in the original paper.

#### Graph annotation growth

1 to 16 annotations were transferred on the graph using GrAnnoT, with the same order as the Panoryza analysis (see above). For all conditions, the number of input genes, graph genes, and graph SG were counted and plotted.

#### Target genome annotation growth

1 to 16 annotations were transferred graph then to Nipponbare using GrAnnoT, with the same order as before (see above). Then, Nipponbare genes were counted for all conditions.

### GrAPEx benchmarks

#### Reduction of the amount of sequence to annotate

All full genomes (excluding Larhamugad) were annotated with Helixer. The graph was then annotated with GrAPEx, with only Larhamugad annotated. These commands were timed using */usr/bin/time*. During GrAPEx’s run, the temporary file that stores the positions to extract was retrieved. The cumulated length of extracted sequences was computed, as well as the cumulated genome length.

#### Increase in the annotated portion of the graph

After annotating the pangenome with Helixer on full genomes or with GrAPEx, the graph was annotated with different annotation sets : the original genomes annotations, or GrAPEx annotation, or Helixer annotation on full genomes. Each time, the number of SG in the graph was retrieved. For GrAPEx’s annotation, the graph annotation was transferred on Nipponbare, and Nipponbare genes were counted.

#### Impact of fragmentation on annotation quality

GrAPEx and Helixer annotations on full genomes were compared on genome ZS97. The file storing the positions of the extracted sequences was retrieved (see above), and used to filter the annotations to compare the annotations of the same regions. Next, *bedtools intersect* was used to compute the overlap between the two annotations, and its output was parsed.

## Acknowledgements

The authors acknowledge the ISO 9001 certified IRD i-Trop HPC (South Green Platform) at IRD Montpellier for providing HPC resources that have contributed to the research results reported within this paper. URL: https://bioinfo.ird.fr/- http://www.southgreen.fr

## Appendix A SynteGroup cluster overlap

**Fig. A1.**
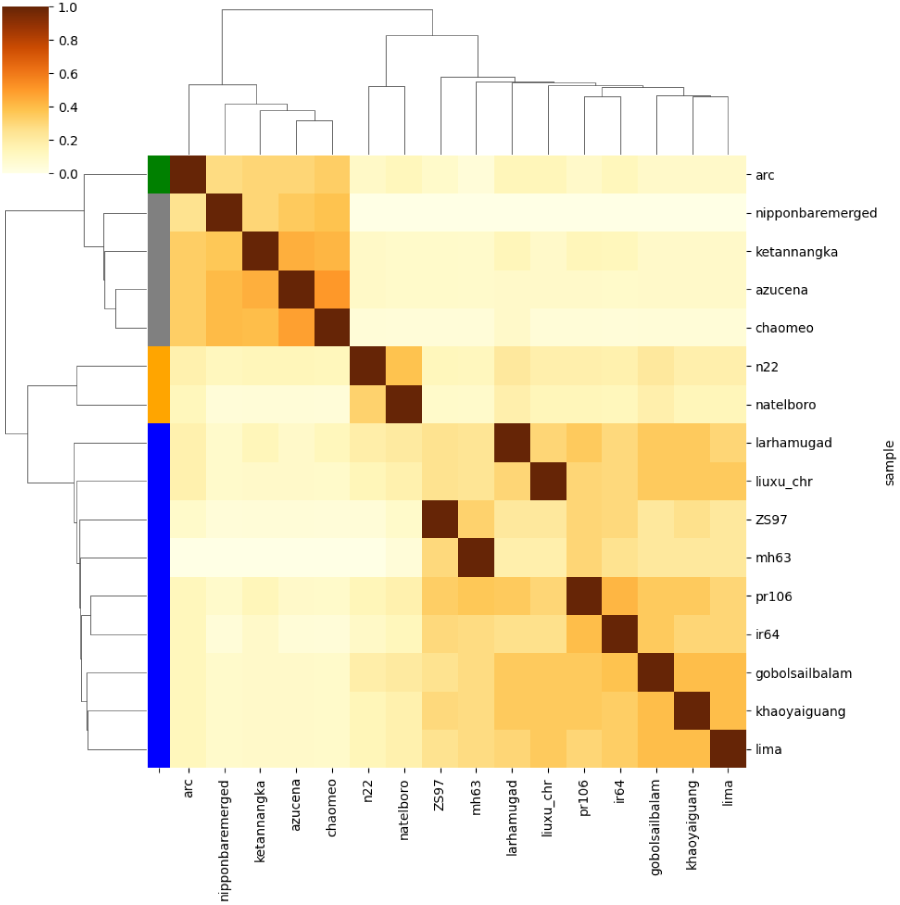
SynteGroup cluster overlap. Matrix of shared SG between genomes, using Nipponbare filtered annotation.

## Appendix B Dispensable gene retrieval

To find if gene SUB1A was annotated with GrAPEx, the graph was annotated with GrAPEx output, and the annotation of genome IR64 was retrieved. As the position of SUB1A is known on IR64, the presence of SUB1A could be attested. The CDS predicted by Helixer on GrAPEx’s fragment corresponds exactly to the expected position of the gene SUB1A.

## Appendix C GrAPEx’s extracted fragments size

**Fig. C2.**
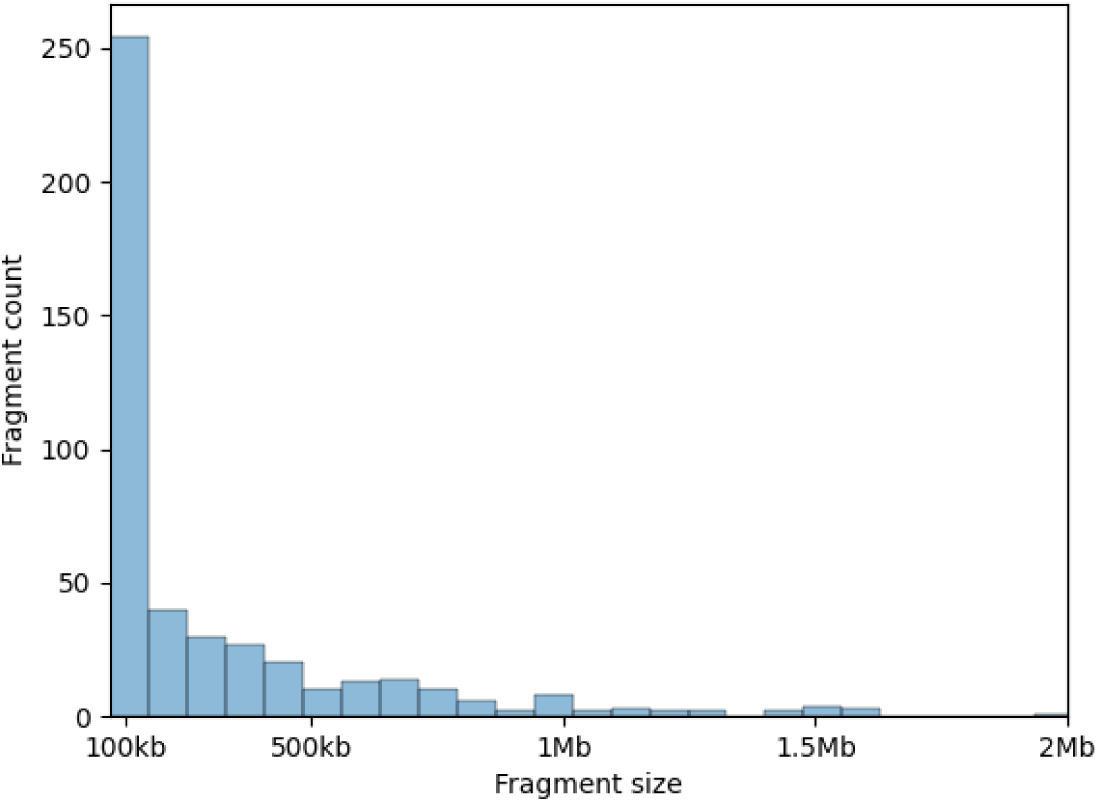
GrAPEx’s extracted fragments size. Most fragments are around 100 kb long. The figure’s x-axis is limited to 2 Mb, but larger fragments exist.

